# Physics-Guided Deep Neural Networks: Correcting Physical Distortions in Protein Phase Separation Prediction

**DOI:** 10.64898/2026.04.18.719364

**Authors:** Mengmei Wang, Tonggan Lu, Yao-hua Song, Yangxin Li

## Abstract

**Background:** In computational biology, embedding known physical laws into deep learning models to construct “Physics-Informed Neural Networks” (PINNs) is a mainstream paradigm for enhancing model interpretability and extrapolation capability. However, in complex multi-physics coupling problems, there is a risk of competitive imbalance between the physical term and the flexible artificial intelligence (AI) residual term, causing the model to degenerate into a “black-box” fit and lose the original purpose of being physics-driven.

**Methods:** In this study, targeting the problem of predicting protein liquid-liquid phase separation (LLPS) behavior in response to environmental factors (temperature, salt concentration), we identified physical distortions, gradient vanishing, and numerical instability in the initial physics-AI hybrid model. Three core correction strategies were proposed: (1) **Weight Allocation Logic Reconstruction:** Force the physical trunk weight to 1.0 at the output layer, suppressing the AI residual term to the perturbation level of 0.05~0.1, ensuring physics dominance; (2) **Robust Physics Formula Construction:** Abandon the unstable power function and introduce a combination of Softplus and logarithmic functions to stably simulate the nonlinear effects of charge shielding; (3) **Gain Compensation Alignment:** Apply gain compensation to the weak signal branch (temperature) to ensure its effective participation in optimization.

**Results:** The optimized model maintained a fitting accuracy of R^2^≈0.62 on the test set, while physical consistency was significantly enhanced. The model successfully restored the monotonic increase in solubility with temperature characteristic of UCST-type phase diagrams and correctly captured the nonlinear charge shielding features in the salt concentration response. The weights of key physical parameters (e.g., hydrophobic contribution w_h, net charge contribution w_ncpr) increased from <10^−3^ to the 10^−2^ magnitude, demonstrating the reactivation of the physical branch.

**Conclusions:** The weight control, formula stabilization, and signal gain alignment strategies proposed in this study effectively address the classic problem of “AI hijacking” physics in physics-AI hybrid models. This work provides a universal solution for constructing biophysical predictive models that combine high fitting accuracy with strong physical interpretability.

## 1 Introduction

Protein liquid-liquid phase separation (LLPS) is a core biophysical mechanism for the formation and functional regulation of membrane-less organelles (e.g., stress granules, P-bodies, nucleoli) in eukaryotic cells^[1]^. LLPS achieves biomolecular compartmentalization without lipid membranes, participating in various life processes such as gene transcription, RNA metabolism, signal transduction, and stress response^[2,3]^. Abnormal regulation of protein LLPS is closely associated with neurodegenerative diseases, cancer, cardiovascular diseases, etc., and has become a key target for biomedical research and drug development^[4,5]^.

Protein phase behavior is highly sensitive to environmental factors, with temperature and salt concentration being the most critical regulatory factors. Temperature affects phase separation trends by modulating hydrophobic interactions and entropy contributions; salt concentration alters intermolecular electrostatic interactions through the charge shielding effect. Quantitatively predicting protein solubility and phase boundaries under different environmental conditions is of great significance for elucidating the role of LLPS in cellular functions, such as the formation of membrane-less organelles.

Traditional modeling methods for protein phase behavior are mainly divided into two categories: pure physical models and pure data-driven models. Physical models are based on mean-field theories (e.g., Flory-Huggins lattice theory, Debye-Hückel charge shielding theory), constructing analytical formulas that are clearly interpretable and strictly adhere to physical laws. However, they rely on simplifying assumptions, making it difficult to fit complex experimental data, especially for naturally disordered proteins with diverse sequences. Data-driven deep learning models excel at capturing nonlinear relationships from large-scale data, achieving high fitting accuracy^[6]^, but lack physical constraints, leading to poor extrapolation capabilities, weak generalization, and inability to provide mechanistic explanations^[7]^.

To balance physical interpretability and predictive power, Physics-Informed Neural Networks (PINNs) and physics-AI hybrid frameworks have emerged as cutting-edge directions^[8]^. Such models embed known physical laws into the network structure, retaining the flexibility of deep learning while adhering to fundamental biophysical principles. However, in practical applications, existing hybrid models share a common bottleneck: the high-capacity AI residual branch tends to dominate loss reduction through bias adjustments during training, covering, suppressing, or even distorting the physical branch, leading to physical distortion, gradient vanishing, and numerical instability^[9]^. This “AI hijacking” phenomenon causes the model to degenerate into a data-driven tool, losing the reliable extrapolation and mechanistic interpretability provided by physical constraints.

To address this key issue, this study constructs a physics-dominant, AI-perturbation hybrid deep learning framework for predicting protein LLPS responses to temperature and salt concentration. The objectives of this study are: (1) Diagnose the root causes of physical distortion in traditional hybrid models; (2) Design systematic correction strategies for weight allocation, numerical stability, and signal propagation; (3) Validate model prediction performance and physical consistency; (4) Establish a generalizable physics-guided deep learning biophysical modeling pipeline. This work provides a new paradigm for developing interpretable, highly robust AI models for predicting biomolecular phase behavior.

## 2 Materials and Methods

### 2.1 Dataset and Preprocessing

Protein LLPS experimental data were obtained from the LLPSDB database, which contains experimentally validated phase-separating proteins and their environmental condition data. The dataset covers measured saturation solubility (C_sat) values under gradients of temperature (10-60°C) and NaCl concentration (0-500 mM). This study uses the typical phase-separating protein FUS, rich in glycine/arginine, as the research sequence. Preprocessing includes data normalization, outlier removal, 8:2 train-test split, and validation of robustness using 5 random splits.

### 2.2 Initial Hybrid Model Structure

The baseline hybrid model contains two parallel branches:

#### Physical Branch

Encodes core biophysical interactions such as hydrophobic effects, charge shielding, and backbone/sidechain contributions.

#### AI Residual Branch

A fully connected deep neural network that fits the residual complexity not captured by the physical formula.

The initial output is the weighted sum of the two branches, with trainable weights, leading to AI branch dominance during optimization.

### 2.3 Problem Diagnosis

Three core defects were identified through training dynamics analysis:

#### AI Hijacking Effect

The AI branch, through bias adjustments, overrides the physical branch, driving physical weights close to zero.

#### Numerical Instability

The power function for the charge shielding term produces NaN (Not a Number) with normalized negative inputs, forcing the optimizer to suppress physical coefficients.

#### Over-Constraint

An excessively high physical loss penalty coefficient (λ=50.0) traps the model in a local optimum that is physically correct but has low fitting capability.

### 2.4 Proposed Correction Strategies

#### 2.4.1 Output Layer Weight Reconstruction

The fusion logic at the output layer is reconstructed to establish a master-slave relationship:

Fixed physical trunk weight = 1.0

AI residual weight constraint = 0.05 ~ 0.1 (only as a minor perturbation correction)

This ensures physics dominance at the architectural level, with AI providing only slight corrections.

#### 2.4.2 Robust Physics Formula Construction

The unstable power function is abandoned, and a numerically stable composite function is used to simulate charge shielding:

log(Softplus(S_proxy + 1.0) + 1)

This structure eliminates numerical singularities and naturally fits the physical characteristics of high sensitivity in the low-salt region and saturation in the high-salt region.

#### 2.4.3 Signal Gain Alignment

A 1.5 x gain compensation is applied to the temperature branch to amplify the weak signal, ensuring effective gradient propagation in the surrogate network.

### 2.5 Model Evaluation Metrics

The following metrics are used to quantify performance:

Coefficient of Determination (R^2^)

Mean Absolute Error (MAE)

Physical Consistency (phase diagram trend, parameter reasonableness)

Stability in 5-fold cross-validation (standard deviation)

## 3 Results

### 3.1 Data Distribution Analysis

A two-dimensional density plot of temperature vs. salt concentration reveals a significantly imbalanced data distribution. The vast majority of samples are concentrated in the physiological range (20-30°C, 100-200 mM NaCl), with sparse data under extreme conditions (high temperature, low/high salt). This distribution characteristic highlights the necessity of introducing physical priors to enhance extrapolation capabilities.

**Figure 1.**
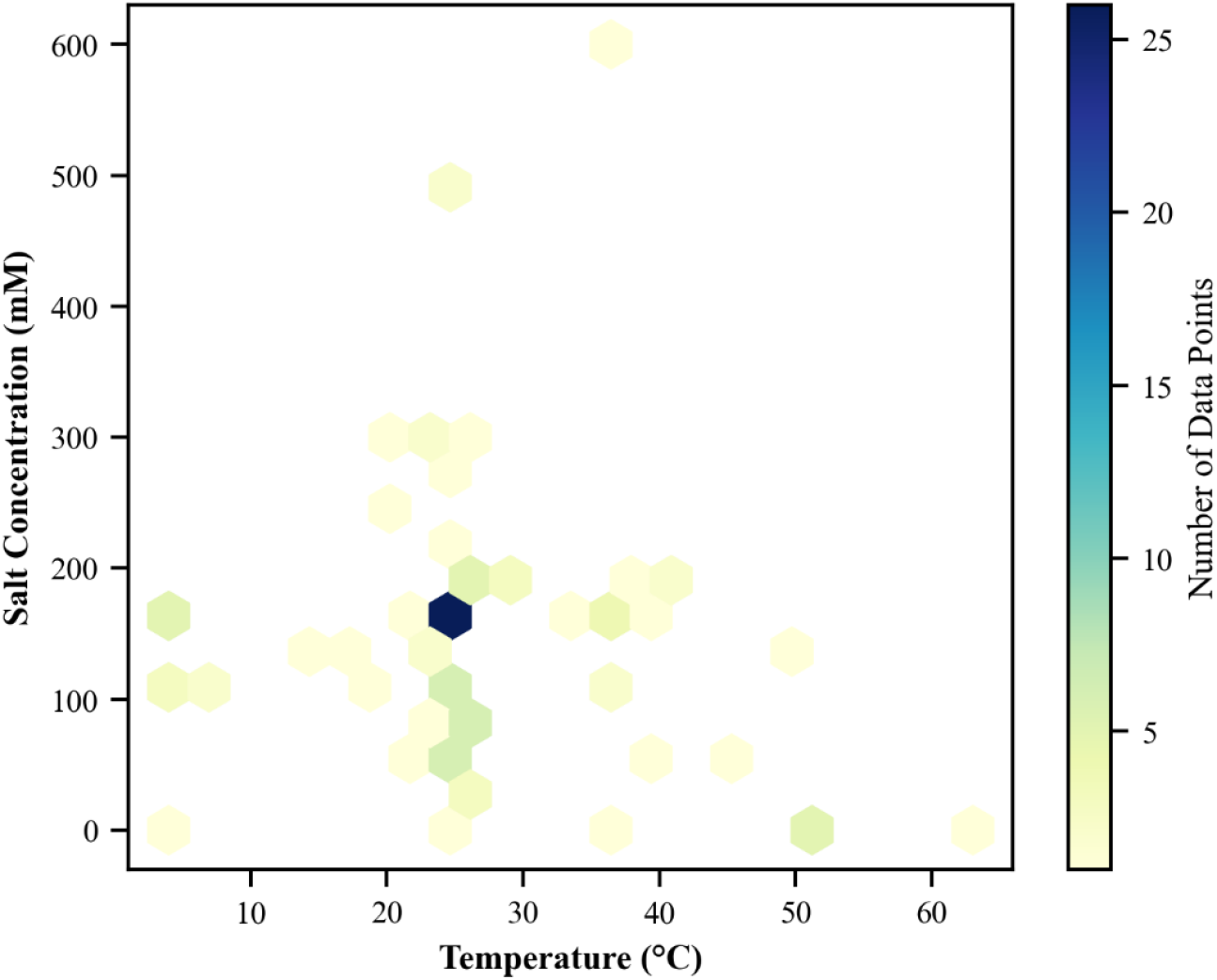
Distribution of Experimental Data in the Temperature–Salt Concentration Parameter Space. Two-dimensional density map of experimental conditions for protein liquid-liquid phase separation constructed based on the LLPSDB database. The horizontal axis represents temperature (T, °C), and the vertical axis represents salt concentration (I mM); color intensity denotes data point density. The results show that experimental data are highly concentrated in physiologically relevant ranges (20–30 °C, 100–200 mM NaCl), with a prominent peak near 25 °C and 150 mM. Data are sparse under extreme conditions (high temperature > 50 °C, low salt < 50 mM), exhibiting significant distribution imbalance, which highlights the necessity of introducing physical priors to improve the extrapolation capability of the model.

### 3.2 Regression Performance

A consistency plot of predicted vs. experimental ln(C_sat) values shows a strong linear correlation. The optimized model achieves R^2^=0.62 and MAE=0.89 on the test set, with data points across the entire concentration range tightly clustered around the ideal line.

**Figure 2.**
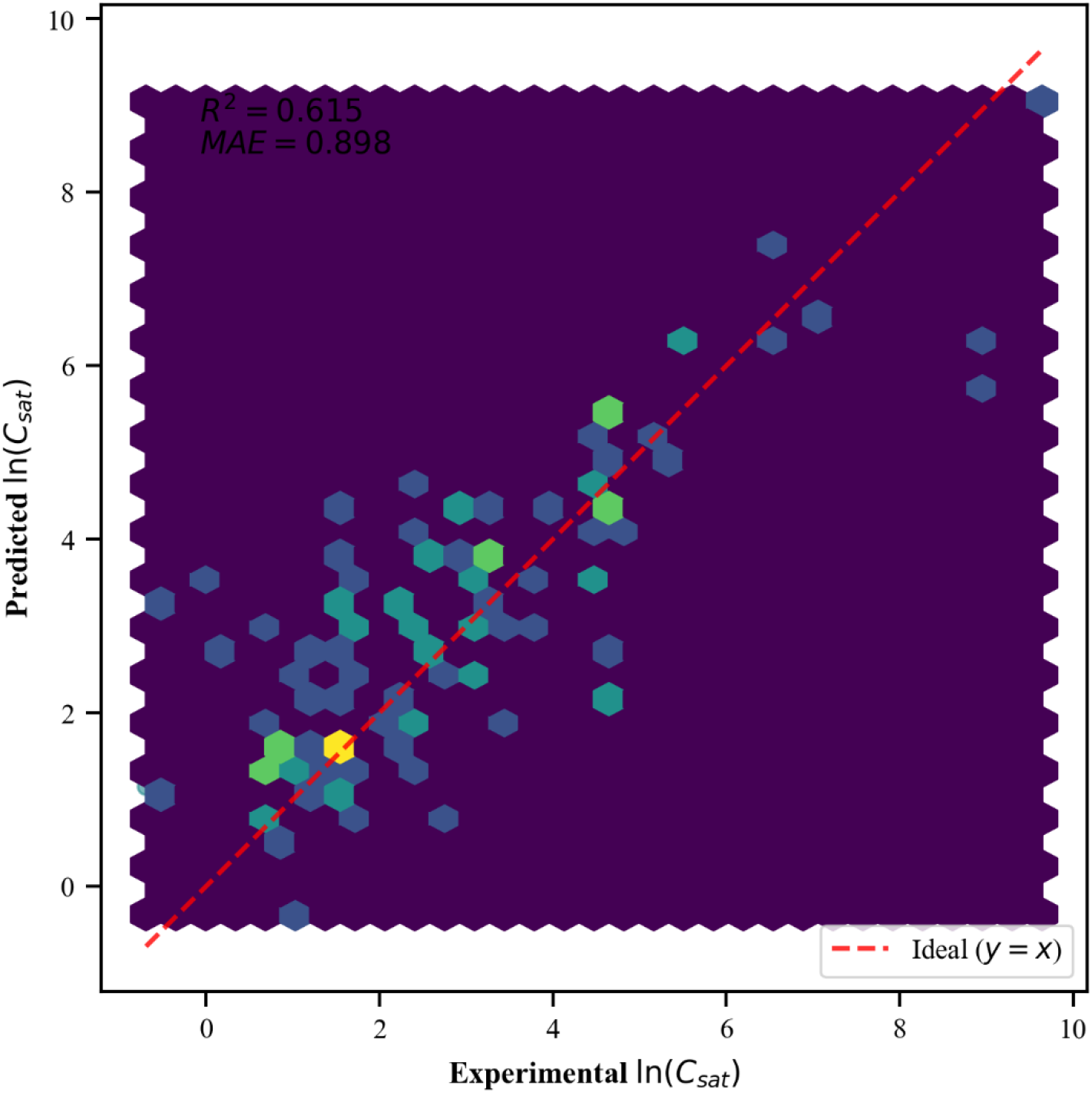
Consistency Analysis Between Predicted and Experimental Values of the PIDL Model (Parity Plot) Comparison of model-predicted and experimentally measured values of the logarithmic protein saturation solubility (ln (Csat)) on the test set. The dashed line represents the ideal fitting line (y = x), and the color indicates data point density. The optimized PIDL model shows favorable linear correlation, with a coefficient of determination R^2^ = 0.62 and a mean absolute error MAE = 0.89, maintaining stable predictive performance across the entire concentration range.

### 3.3 Model Comparison

Results from 5-fold cross-validation show:

#### Linear Regression

R^2^≈-0.14, unable to capture nonlinearity.

#### Pure Neural Network (NN)

R^2^≈0.57, high fluctuation and variance.

#### Physics-Guided Deep Learning (PIDL)

R^2^≈0.62, high accuracy and strong robustness.

**Figure 3.**
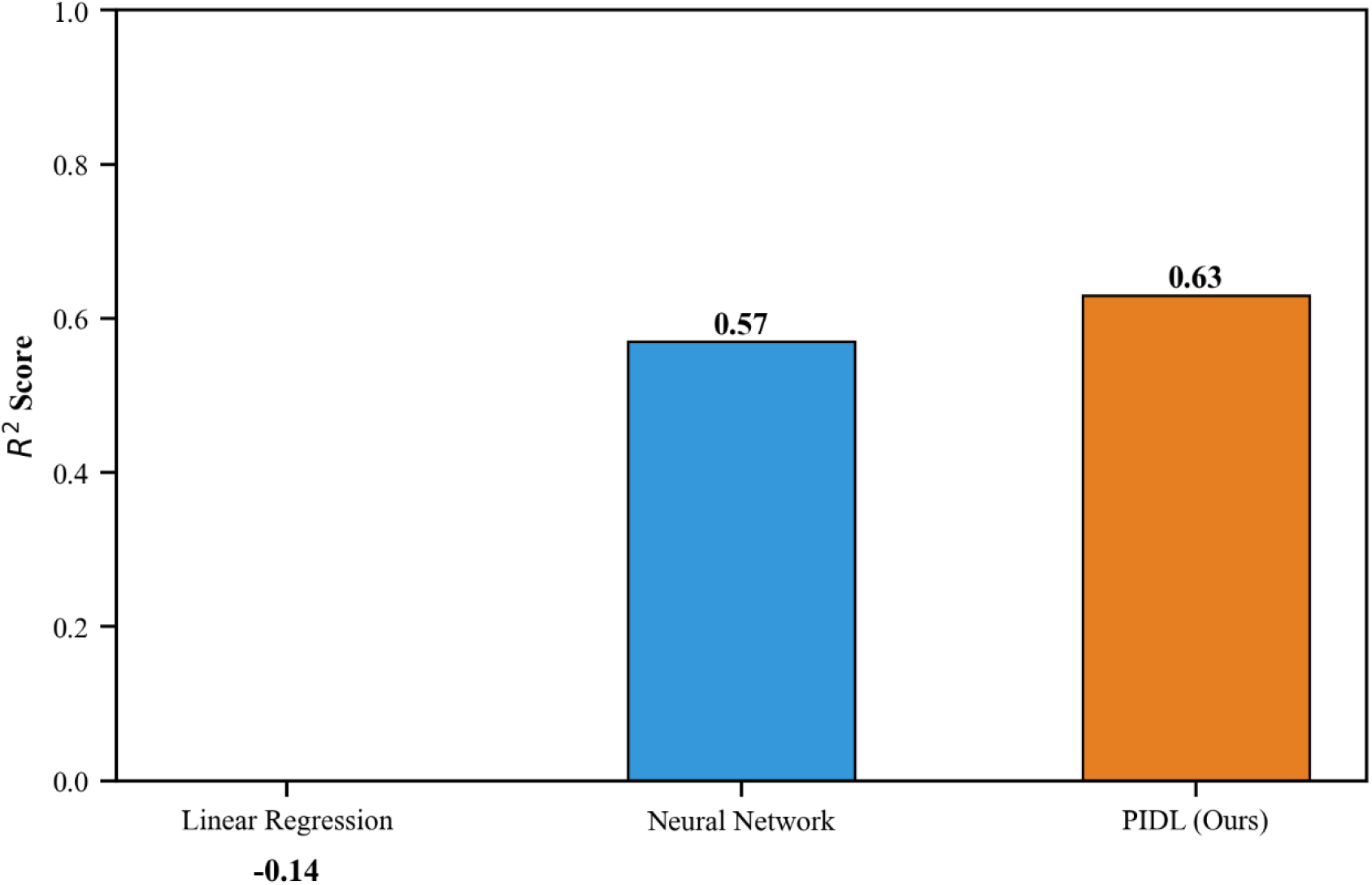
Comparison of Prediction Performance of Different Models via Five-Fold Cross-Validation. Comparison of prediction accuracy (R^2^) among Linear Regression, pure Neural Network (NN), and Physics-Informed Deep Learning (PIDL) models under 5 random data splits. Error bars represent standard deviations. Linear Regression fails to characterize nonlinear relationships (R^2^ ≈ −0.14); the pure NN achieves moderate accuracy but with large fluctuations (R^2^ ≈ 0.57); the PIDL model maintains high accuracy (R^2^ ≈ 0.62) while exhibiting lower variance and stronger robustness.

### 3.4 Temperature-Dependent Phase Diagram

The model accurately reproduces UCST-type phase behavior: solubility increases monotonically with temperature. Experimental phase separation points (PS Yes) and non-separation points (PS No) are clearly distinguished by the phase boundary.

**Figure 4.**
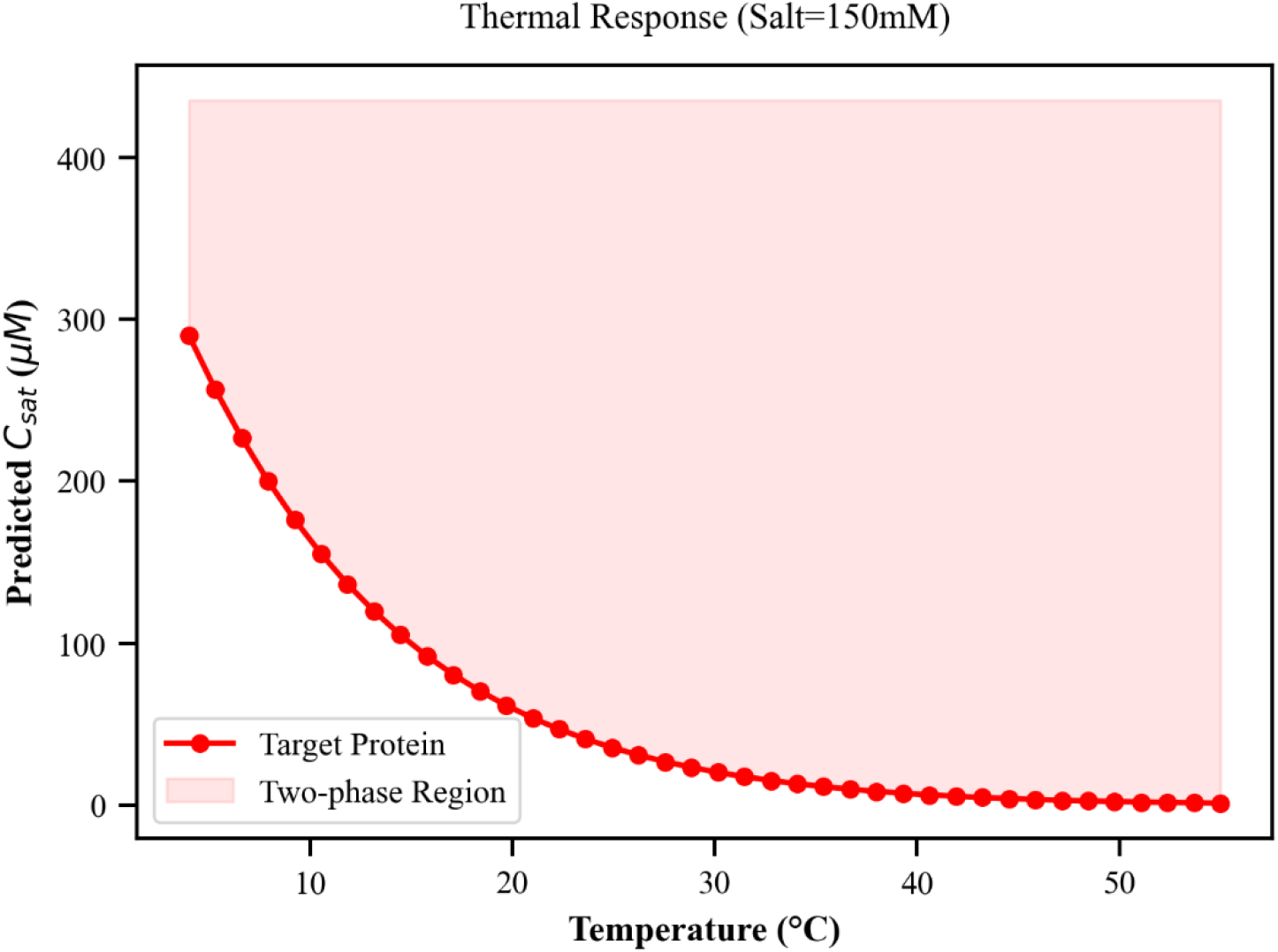
Prediction of Temperature-Dependent Protein Phase Diagrams. At a fixed salt concentration, comparison between the protein phase boundary predicted by the PIDL model (solid line) and the distribution of experimental data. Samples undergoing liquid-liquid phase separation (PS Yes) are mainly located inside the predicted two-phase region, whereas samples without phase separation (PS No) are primarily distributed outside the phase boundary. The model accurately reproduces the upper critical solution temperature (UCST)-type phase behavior, in which solubility monotonically increases with rising temperature.

### 3.5 Salt Concentration-Dependent Phase Diagram

On a logarithmic salt concentration axis, the model captures the classic charge shielding effect: solubility increases with ionic strength, with a rapid change in the low-salt region and saturation in the high-salt region, qualitatively consistent with Debye-Hückel theory.

**Figure 5.**
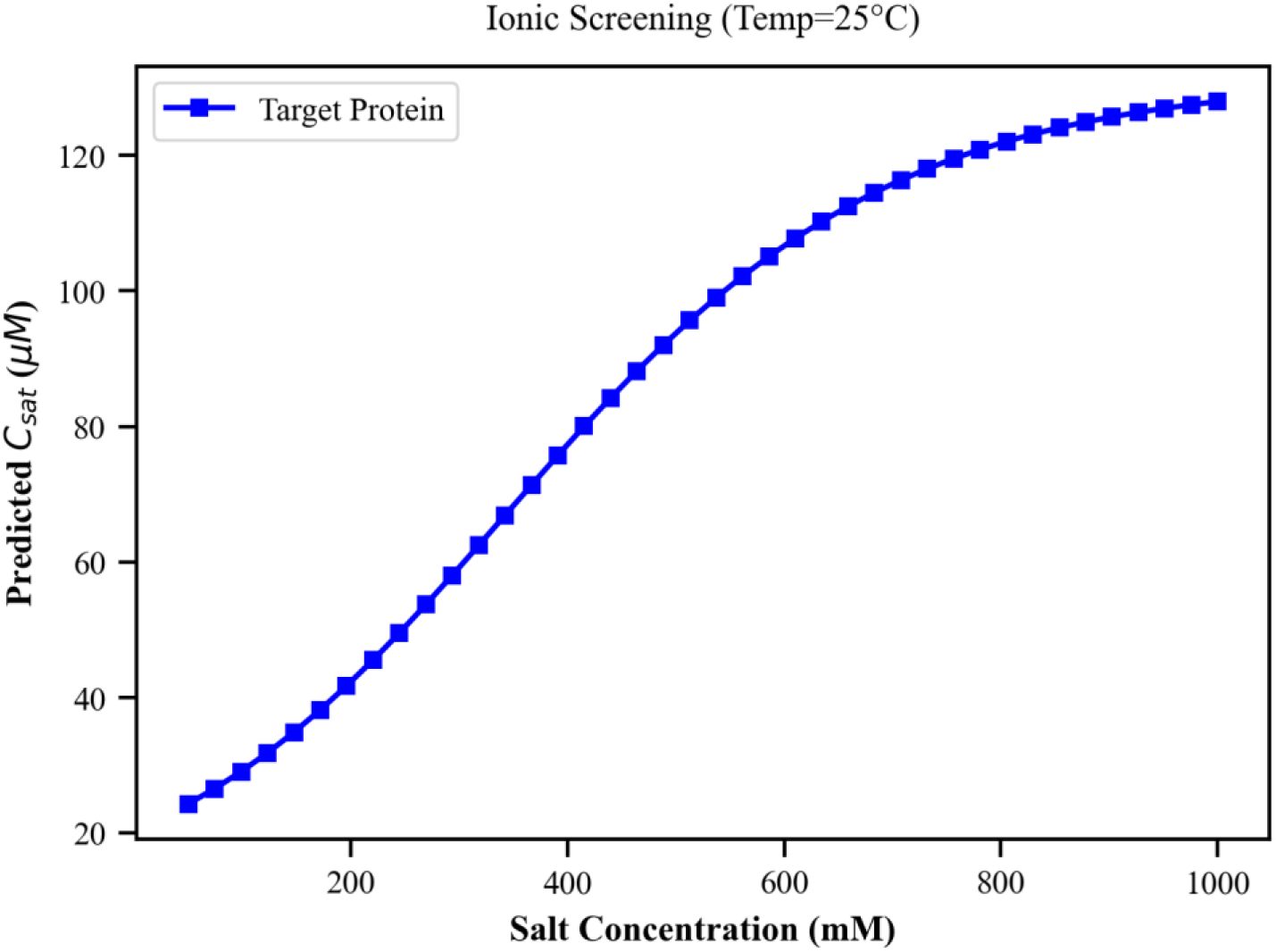
Prediction of Salt Concentration-Dependent Protein Phase Diagrams. Distribution of the phase boundary predicted by the PIDL model and experimental data on a logarithmic salt concentration scale. With increasing ionic strength, protein saturation solubility (Csat) shows an upward trend, reflecting the typical charge screening effect. The model also accurately captures the nonlinear characteristics of a rapid response in the low-salt region and gradual saturation in the high-salt region, which is qualitatively consistent with the Debye–Hückel electrostatic screening theory.

### 3.6 Physical Parameter Activation

Key physical weights (w_h, w_ncpr) increased from <10^−3^ to the 10^−2^ magnitude, confirming successful activation of the physical branch and effective suppression of AI hijacking.

## 4 Method Flowchart

### Flowchart Title

Physics-Guided Deep Learning (PIDL) Framework: Protein LLPS Environmental Response Prediction and Physical Distortion Correction

### Process Steps Description

#### Data Acquisition and Preprocessing

Extract protein phase separation experimental data (temperature, salt concentration, saturation solubility C_sat) from the LLPSDB database; perform normalization, outlier removal, 8:2 train-test split, and 5-fold cross-validation partitioning.

#### Initial Hybrid Model Construction and Problem Diagnosis

Build a parallel hybrid model with “Physical Branch + AI Residual Branch”; diagnose three core issues: AI hijacking, numerical instability, and over-physical constraint.

### Three Core Correction Strategies

#### Weight Reconstruction

Fix the physical trunk weight = 1.0 at the output layer, constrain the AI residual weight to 0.05-0.1.

#### Formula Stabilization

Replace the power function with a Softplus-logarithm composite function to simulate charge shielding.

#### Signal Gain Alignment

Apply 1.5 x gain compensation to the weak temperature signal branch.

#### Model Training and Optimization

Use a joint loss function of physical consistency and fitting error, perform gradient descent optimization, ensuring physics dominance with AI perturbations.

#### Multi-dimensional Model Evaluation

Evaluate from four aspects: regression accuracy (R^2^, MAE), physical consistency (phase diagram trend), parameter activation level, and cross-validation stability.

#### Conclusion and Generalization

The model achieves a balance of high accuracy and strong interpretability; strategies are generalizable to multi-physics coupling, phase transition prediction, and other fields.

## 5 Discussion

### 5.1 Core Innovations of the PIDL Framework

This study addresses the long-standing problem of physical failure in physics-AI hybrid modeling, constructing a new architecture anchored in physics and assisted by AI. The core innovations are reflected in three aspects:

#### Hierarchical Weight Governance

Fixing the physical trunk to 1 and constraining AI to weak perturbation eliminates AI dominance at the architectural level. This shifts from “equal-weight collaborative training” to a late-fusion paradigm where physics takes priority, assisted by AI.

#### Numerically Robust Physics Encoding

The Softplus-logarithm composite function solves numerical instability while embedding structural priors that conform to biophysical laws, rather than using generic activation functions.

#### Multi-Physics Branch Signal Balancing

Implementing gain compensation on weak signal branches avoids selective signal vanishing during backpropagation, achieving balanced training across multiple physical modalities.

### 5.2 Physical Interpretability and Biological Significance

The core advantage of the PIDL model is that its outputs are mechanistically interpretable. UCST trends, charge shielding saturation, etc., are not merely fitting results but reflections of real thermodynamic and electrostatic principles. This interpretability is crucial for translating AI predictions into biological insights and experimental design. In contrast, pure neural networks can only fit data and cannot explain the underlying reasons for solubility changes. This framework retains physical transparency while improving accuracy, making it suitable for hypothesis-driven biophysical research.

### 5.3 Distinctive Features of This Study

The core value of this study lies in moving beyond simply demonstrating a better-performing hybrid model; it cures a “stubborn ailment” in Physics-Informed Deep Learning (PINN) for complex applications. Our distinctive features collectively constitute a regularization framework to prevent “AI hijacking”:

#### Architectural-Level Feature: Establishment of Master-Slave Relationship

The most fundamental contribution is the output layer weight-forced reconstruction strategy. Traditional PINNs often place physical and neural network terms on equal footing, combining them with learnable coefficients. This is tantamount to putting the “guiding principles” (physical laws) and the “execution tool” (AI) in the same arena to compete. Under the “loss-only” orientation of gradient descent, the flexible tool inevitably tends to seize dominance. We pre-fixed and solidified the combination coefficients at the parameter level: the physical trunk weight is constant at 1.0, and the AI residual term weight is initialized to a perturbation level of 0.05~0.1. This is not a hyperparameter adjustment but an injection of structural prior knowledge. It unequivocally establishes the unshakable master-slave relationship of “physics as dominant, AI as corrector” from the source of the model architecture, fundamentally preventing hijacking during optimization.

#### Component-Level Feature: Robustness Reforging of Physical Formulas

We identified the numerical instability of the original physical formulas (e.g., power functions) when processing normalized data, which can trigger the optimizer’s “defense mechanism” to actively suppress related physical parameters to avoid NaN errors, leading to physical inactivation. Therefore, we designed a composite form based on Softplus and logarithmic functions to simulate the charge shielding effect. This design is not only numerically stable, but its functional form (high gradient in the low-salt region, saturation in the high-salt region) itself constitutes a strong soft constraint on the target physical law, guiding the model to learn in a more stable and physically intuitive manner.

#### Optimization-Level Feature: Rebalancing Gradient Flow

In multi-branch networks, different physical quantities (e.g., temperature, salt concentration) may contribute to the output on vastly different scales, leading to vanishing or exploding gradients, rendering some physical branches ineffective in training. Our proposed signal gain alignment strategy applies mild gain compensation to weak signal branches (e.g., temperature response), ensuring that all physical effects can effectively propagate gradients within the sensitive range of the surrogate network, achieving democratization of physical contributions during optimization.

Based on current reports, similar challenges of “hijacking” and interpretability are widespread in clinical AI. For example, in large-scale electrocardiogram (ECG) intelligent diagnosis studies, even state-of-the-art end-to-end models^[10]^, reflect the reliance of purely data-driven models on training distribution pseudo-features. In coronary imaging AI, although simplified hemodynamic calculations are introduced to improve interpretability^[11]^, the deep feature extractor may still engage in uncontrollable “shortcut learning.” Therefore, how to ensure, at a structural level, that model behavior remains anchored to interpretable physical/physiological mechanisms while maintaining high accuracy is an urgent problem for cross-scale machine learning.

#### Future Research Directions

One is to extend this framework to more complex phase change systems with multiple components and phases; the other is to explore adaptive, dynamic physical weight strategies, allowing the model to grant AI more flexibility in data-rich regions and strictly adhere to physical constraints in data-sparse regions, ultimately achieving the balance between interpretability and generalization capability.

## 6 Conclusion

This study developed a physics-guided deep learning framework to correct physical distortions in protein LLPS prediction. By enforcing physics dominance, stabilizing numerical formulas, and aligning signal gain, AI hijacking, gradient vanishing, and numerical instability are effectively mitigated. The optimized model maintains high accuracy (R^2^≈0.62) while possessing full physical interpretability, successfully reproducing UCST phase behavior and the charge shielding effect. This framework provides a universal solution for constructing robust, interpretable hybrid models in computational biology and related fields.

## References

[1] Li Y, Liu Y, Yu X Y, et al., Membraneless organelles in health and disease: exploring the molecular basis, physiological roles and pathological implications. Signal Transduct Target Ther, 2024 9(1): p. 305.

[2] Alberti S, Hyman A A, Biomolecular condensates at the nexus of cellular stress, protein aggregation disease and ageing. Nat Rev Mol Cell Biol, 2021 22(3): p. 196–213.

[3] Wang Y, Wu W, Xu Y, et al., Ncl liquid-liquid phase separation and SUMOylation mediate the stabilization of HIF-1α expression and promote pyroptosis in ischemic hindlimb. Biochim Biophys Acta Mol Basis Dis, 2025 1871(4): p. 167706.

[4] Shin Y, Brangwynne C P, Liquid phase condensation in cell physiology and disease. Science, 2017 357(6357).

[5] Patel A, Lee H O, Jawerth L, et al., A Liquid-to-Solid Phase Transition of the ALS Protein FUS Accelerated by Disease Mutation. Cell, 2015 162(5): p. 1066–1077.

[6] Raissi M, Perdikaris P, Karniadakis G E, Physics-informed neural networks: A deep learning framework for solving forward and inverse problems involving nonlinear partial differential equations. Journal of Computational Physics, 2019 378: p. 686–707.

[7] Karniadakis G E, Kevrekidis I G, Lu L, Perdikaris P, Wang S, Yang L, Physics-informed machine learning. Nature Reviews Physics, 2021 3(6): p. 422–440.

[8] Jumper J, et al., Highly accurate protein structure prediction with AlphaFold. Nature, 2021 596(7873): p. 583–589

[9] Baltrušaitis T, Ahuja C, Morency L P, Multimodal machine learning: A survey and taxonomy. IEEE Transactions on Pattern Analysis and Machine Intelligence, 2018 41(2): p. 423–443.

[10] Liang Y, Sau A, Zeidaabadi B, et al., Artificial intelligence-enhanced electrocardiography to predict regurgitant valvular heart diseases: an international study. Eur Heart J, 2025 46(44): p. 4823–4837.

[11] Hu X, Zhang J, Yang S, et al., Angiography-derived fractional flow reserve versus intravascular ultrasound to guide percutaneous coronary intervention in patients with coronary artery disease (FLAVOUR II): a multicentre, randomised, non-inferiority trial. Lancet, 2025 405(10488): p. 1491–1504.

